# Macrophage activation drives ovarian failure and masculinization

**DOI:** 10.1101/2023.01.03.522645

**Authors:** Paloma Bravo, Yulong Liu, Bruce W. Draper, Florence L. Marlow

## Abstract

In humans, premature ovarian insufficiency (POI) is caused by autoimmunity and genetic factors, such as mutation of BMP15, a key ovarian determining gene. The cellular mechanisms associated with ovarian failure caused by BMP15 mutation and immune contributions to the disorder are not understood. BMP15’s role in ovarian follicle development is conserved in vertebrates, including zebrafish. Using zebrafish, we established a causal link between macrophage activation and ovarian failure. We identified a germline-somatic gonadal cell-macrophage axis underlying ovarian atresia. Germline loss of Bmp15 triggers this axis that single-cell RNA sequencing and genetic analyses indicate involves activation of ovarian somatic cells that express conserved macrophage-activating ligands. Genetic ablation of macrophages blocks premature oocyte loss. Thus, the axis identified here represents potential therapeutic targets to preserve female fertility.

**One-Sentence Summary:** Sex reversal due to Bmp15 deficiency requires macrophage activation by Csf1a, which is expressed by specialized pre-follicle cells in zebrafish.

## Main Text

In many animals, including mammals, embryos initially develop with indeterminant gonads that in males are remodeled into testis. As in mammals, the zebrafish ovary is comprised of germline cells and their supporting follicle cells of the somatic gonad – granulosa and theca cells – that support differentiation, growth, and maintenance of the germline. In humans, *BMP15* was the first identified ovarian determining gene on the X-chromosome (*1*). BMP15, a conserved ligand produced by early oocytes of most vertebrates, is an essential regulator of follicular growth that in mammals, promotes granulosa cell proliferation (*2-4*). BMP15 secreted by oocytes binds and signals through BMPR2 and SMAD1 to activate targets in the somatic gonad. In mouse, ovarian follicles with high BMP15 expression progress while those with low levels undergo atresia (*5*). Accordingly, BMP15 has been implicated in the pathophysiology of premature ovarian failure (POF) (*6, 7*), which can involve a combination of genetic, endocrine, immune, and environmental factors. Mutation of BMP15 is a predominant genetic cause of premature ovarian insufficiency (POI), a reproductive disorder caused by genetics and immunity-related disorders that affects oocyte quality and leads to hyperandrogenism and inflammation. POI and POF not only lead to sterility, but more broadly affect female health, including bone, cardiovascular and neurological symptoms (*8*). In zebrafish, Bmp15 is also essential for maintenance of the female germline. *bmp15* mutants develop as females but undergo ovarian failure and sex reversal due to a mechanism presumed to involve granulosa cell and estrogen deficiencies (*9*). We recently showed that loss of *bmp15* mutant oocytes could be attenuated by eliminating the conserved masculinizing factor Double sex and Mab3 related transcription factor, Dmrt1 (*10*). However, the mechanism underlying ovarian failure is not understood.

To determine if Bmp15 promotes follicle survival, we generated double mutants (DM) lacking both *bmp15* and tumor suppressor factors (TSF) *tp53* or *chek2*, which can suppress cell death and oocyte loss in some zebrafish and mouse fertility mutants (*11-13*). We found that while *bmp15* heterozygotes and *chek2* or *tp53* single mutants showed no sex bias by 85 days post fertilization (d), all *bmp15* homozygous mutants were fertile males (Fig. 1, A, B and G, S1). Similarly, *bmp15* mutants heterozygous for TSFs were all male by 95d. However, *bmp15;chek2DM*s, but not *bmp15;tp53DM* retained oocytes and female sex traits (Fig. 1, C to G). Significantly, scRNA-seq data indicate that *chek2* is expressed predominantly in oocytes (Fig. S2). Therefore, we conclude that sex reversal of *bmp15* mutants can be suppressed by preventing *chek2-*mediated cell death of oocytes. However, additional somatic fate deficits likely contribute to infertility as mutant follicles did not progress in development even when oocyte loss and sex reversal were blocked. Therefore, Bmp15 has a conserved function in signaling from the oocyte to promote growth and survival of ovarian follicles, which prevents ovarian failure in zebrafish.

**Fig. 1.**
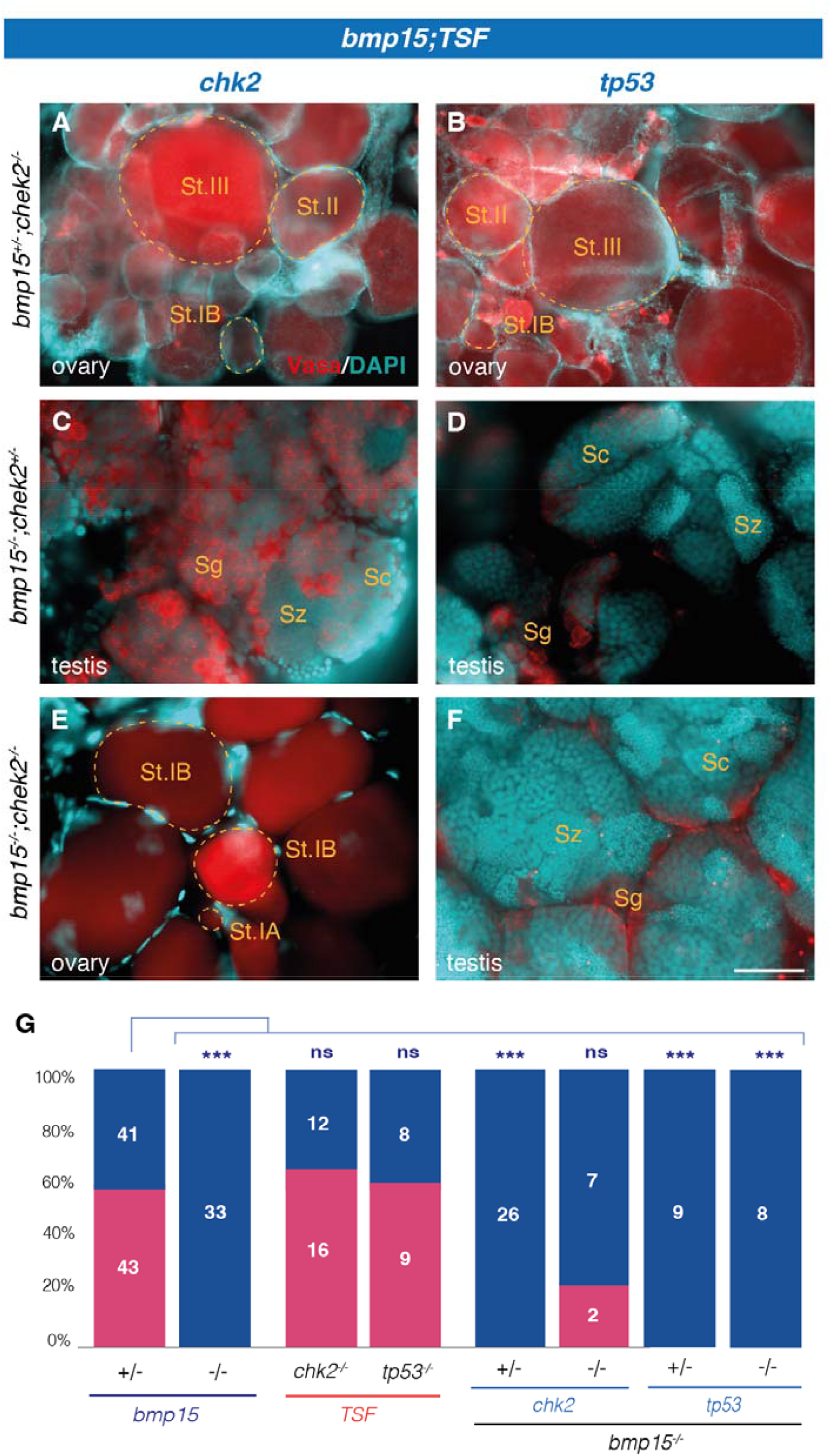
Loss of Chk2 suppresses oocyte loss and masculinization in the absence of Bmp15. **(A-F)** Immunostained adult (A, C, E) *bmp15;chk2* and (B, D, F) *bmp15;tp53* gonads of indicated genotypes. Vasa (red) labels germ cells. **(G)** Adult sex ratios. Female (pink), male (blue). Numbers indicate individuals examined. Statistical analysis: Chi-square test with Bonferroni correction; p-Value comparisons to *bmp15*^+/-^, ***P ≤ 0.0001. Scale bar: (A, B) 250µm, (C-F) 50µm. St.I: stage I oocyte, St.II: stage II oocyte, Sg: spermatogonia, Sc: spermatocyte, Sz: spermatozoa.

Chk2 has been identified as a transcriptional activator of proinflammatory cytokines as well as Tp53-dependent and independent apoptosis (*14*). Given Chk2’s role in proinflammatory cytokine regulation and that immune-related disorders and inflammation are implicated in the pathophysiology of ovarian failure in POF, POI, we investigated immune cell involvement in ovarian failure (*15, 16*). Macrophages are key among innate immune cells in eliminating foreign or dying cells from tissues, regulating their uptake and removal, producing cytokines and factors that recruit and regulate other immune cells, and expressing growth factors to control proliferation, vascularization, and depositing extracellular matrix during wound healing and repair thus contributing to tissue homeostasis (*17-19*). In mammalian ovary, macrophages are abundant and heterogeneous, showing distinct and ovarian cycle dependent distributions and activation states (*20, 21*), and are hypothesized to degrade the follicle by promoting granulosa cell apoptosis (*22-25*). To investigate if macrophages contribute to follicle atresia and masculinization, we genetically ablated macrophages in *bmp15* mutants using two genetic models. In zebrafish, macrophages arise in two waves of specification and colonization. Loss of *colony stimulating factor 1 receptor a (csfr1a)* ablates primitive macrophages, which arise from the anterior lateral plate mesoderm. By contrast, *csf1 receptor b (csf1rb)* mutation affects the definitive population that originate from the ventral dorsal aorta at 14d (*26, 27*). Despite distinct contributions to the primitive and definitive waves, both *csf1ra* and *csf1rb* single mutants have macrophage deficits as adults and DM fish lacking both receptors (*csf1r*^*DM*^) show a lifelong absence of macrophages without affecting sex ratios (Fig. S1A) (*28-30*). Similarly, mutation of *interferon transcription factor 8* (*irf8)*, a conserved regulator of macrophage differentiation ablates macrophages without disrupting normal sexual differentiation (Fig. S1A) (*31-33*). To study macrophage contributions to follicle atresia and sex reversal, we generated *bmp15* mutants lacking all macrophages (Mφ^-^): *csf1r*^*DM*^ or *irf8* mutants, or with deficiency of primitive and/or definitive wave macrophages (Mφ^haploinsufficiency^), *e*.*g*. lacking one *csf1* receptor and heterozygous for the other or *irf8* (Fig. 2 and Fig S1). Analyses of adult gonads revealed that *bmp15* mutants with macrophages were all fertile males while those lacking macrophages retained ovaries but were sterile because *bmp15* mutant oocytes arrest development at an early stage of oogenesis (Fig. 2, A-F and M, and Fig. S1, C-E). Interestingly, double heterozygosity for both *csf1* receptors (*csf1r*^*DH*^) or for *irf8* prevented oocyte loss and sex reversal (Fig S1, B, F-G). Additionally, *csf1ra* mutants heterozygous for *csf1rb*, which eliminates primitive and most definitive macrophages (*29*) prevented ovarian failure and sex reversal (Fig. 2D, and G-M), indicating a threshold number or specific macrophage population mediates sex reversal.

**Fig. 2.**
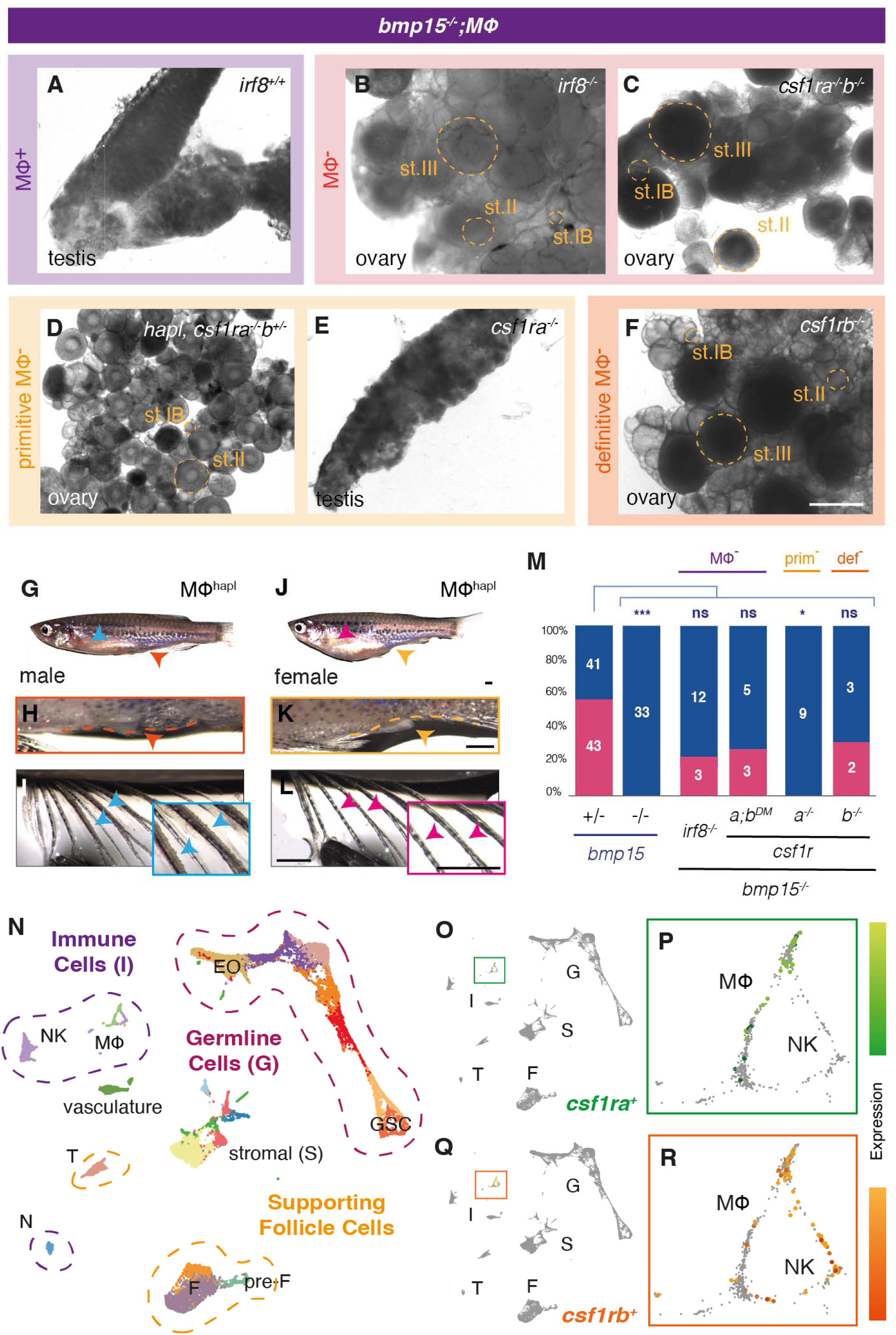
Definitive macrophages are required for ovarian failure and masculinization of *bmp15* mutants. **(A-F)** Live tissue pictures of adult gonads of *bmp15* mutants when macrophages are (A) present, (B, C) completely ablated, (D) haploinsufficient, or lacking (E) primitive, or (F) definitive populations. **(G-L)** Secondary sex traits of adult: (G) male body shape, (H) papilla (orange dashed lines and arrow), and (I) lateral fin tubercles (blue arrows and box); (J) rounded female body shape, (K) distinct papilla (yellow dashed lines and arrow), and (L) absence of tubercles (pink arrows and box). **(M)** Sex ratios of indicated genotypes. Female, pink; male, blue. Numbers indicate individuals examined. Statistical analysis: Chi-square test with Bonferroni correction; p-Value comparisons to *bmp15*^+/-^, *P ≤ 0.0125, ***P ≤ 0.0001. **(N)** UMAP plot of cell types in 40d ovary. **(O-R)** Expression of indicated genes in the ovary in (O, Q) all cells and (P, R) macrophages specifically. N: neutrophils, MΦ: macrophages, NK: natural killer cells, F: follicle, T: theca, EO: early oocyte, GSC: germ stem cell. Scale bar (A-F): 500µm, (G-L): 1mm. St.I: stage I oocyte, St.II: stage II oocyte.

Single-cell RNA sequencing (scRNA-seq) of 40d ovary confirmed the presence of *csf1ra/b*-expressing macrophages in the ovary (Fig. 2, N-R) that also expressed *irf8* (Fig. 3A and E) (*34*). To determine if ovarian failure and sex reversal requires a threshold number or a unique subtype of macrophages, we analyzed compound mutants for various combinations of *csf1ra/b* and found that *bmp15* mutants lacking only *csf1ra* were all male as adults (Fig. 2, E and M). Thus, primitive macrophages are dispensable for ovarian failure and sex reversal. In contrast, loss or haploinsufficiency of definitive macrophages (Fig. 2, F and M and Fig S1B) suppressed ovarian failure and sex reversal. Although *irf8* was also expressed in stromal cells and some pre-follicle cells (Fig. 3, A, C-E), based on the genetic evidence that loss of *csfr1a/b*, which are only expressed in macrophages, or loss of *irf8* both block sex reversal, we conclude that macrophages are direct cellular mediators of sex reversal in the absence of Bmp15. Furthermore, sex reversal requires an activity unique to a specific state or subpopulation of definitive macrophages.

**Fig. 3.**
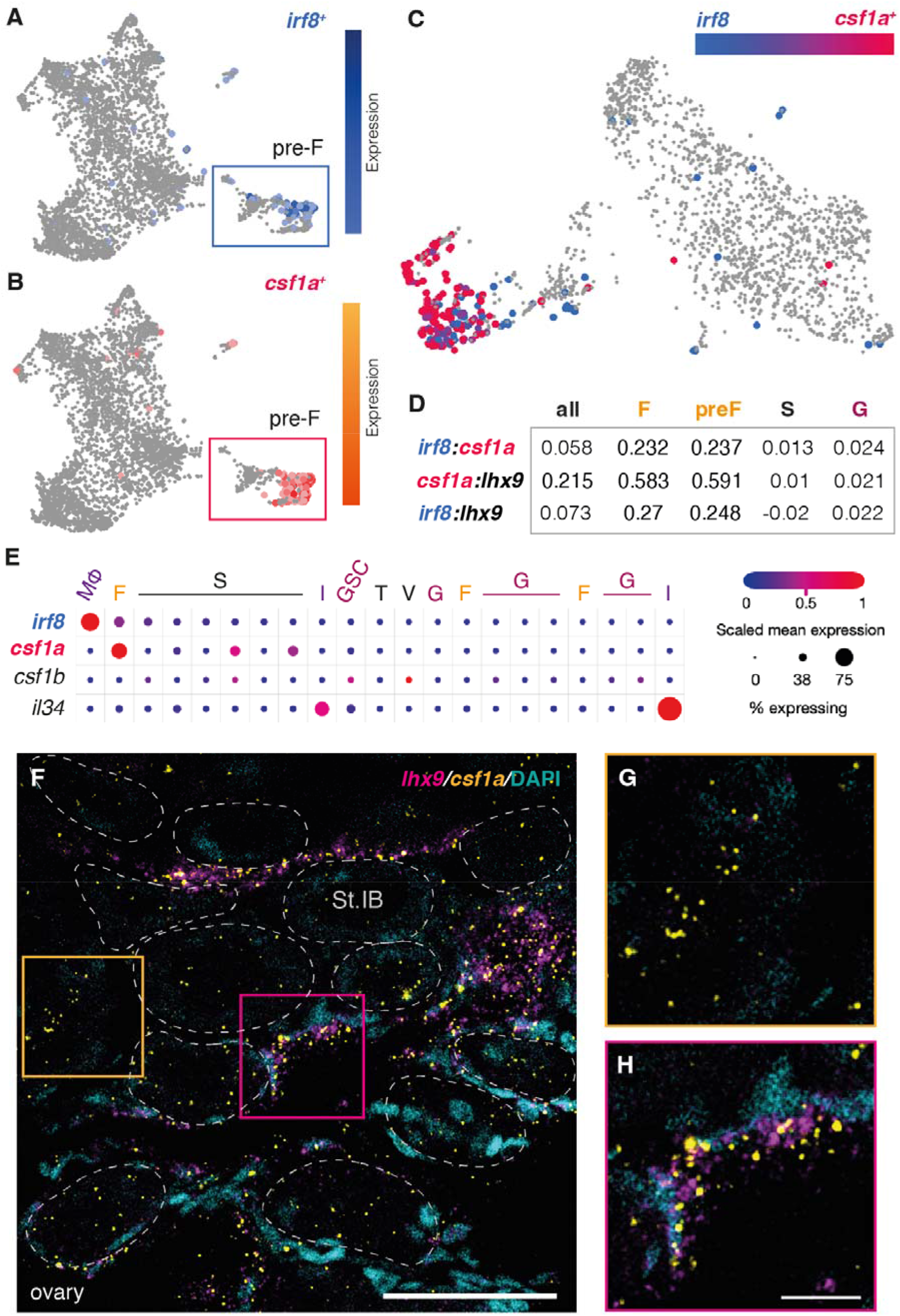
*irf8* and *csf1a* are coexpressed in a subpopulation of pre-follicle cells in the early ovary. **(A-C)** UMAP plot of indicated gene’s expression in the 40d ovary in (A, B) follicle cells, and (C) magnified subpopulation of pre-follicle cells. **(D-E)** Analysis of expression profiles of indicated genes in specified clusters of ovarian cells represented by (D) Spearman’s Rho correlation analysis, and (E) dot plot graph. **(F-H)** Double HCR RNA-FISH confocal images of follicle cells (*lhx9*, magenta) and Csf1rs’ main ligand (*csf1a*, yellow) and DAPI (DNA, blue) in 40d ovary. Regions boxed in (F) show magnified views of (G) *lhx9*-expressing pre-F cells and (H) *lhx9/csf1a* co-expressing MAFCs. F: follicle, G: germline, GSC: germ stem cells, I: immune, MΦ: macrophages, S: stroma, T: theca, V: vasculature.

Csf1 receptors on macrophages are activated by Csf1 ligands and Il-34 (*35, 36*); therefore, the cells that express these ligands represent candidate triggers of sex reversal. Thus, we determined which cells in the ovary express Csf1R activating ligands. scRNA-seq analyses revealed a unique population of ovarian pre-follicle cells that express *lhx9, irf8*, and *csf1a* ligand (Fig. 3B) (*34*). Among known *csf1r* ligands, scRNA-seq indicated that *csf1a* was expressed in pre-follicle cells, whereas *csf1b* was not appreciably expressed and *il34* was restricted to other cell types (Fig. 3, B-E, and Fig. S3). Based on its limited expression we reasoned that Csf1b was not likely the relevant ligand. Although expressed, Il34 is also not likely the key activating ligand since it signals primarily through Csf1Ra, which is dispensable for ovarian failure and sex reversal on its own (Fig. 2E and M). Using fluorescence *in situ* hybridization to verify co-expression of *lhx9* and *csf1a* in 40d wild-type pre-follicle cells, we found that, as indicated by the scRNA-seq data, *csf1a* is expressed in *lhx9* expressing pre-follicle cells and in *lhx9-*negative cells that are likely stromal cells. (Fig. 3F). Thus, we hypothesize a novel role for the subpopulation of pre-follicle cells, hereafter called Macrophage-Activating Follicle Cells (MAFCs), that express *csf1a*, the main ligand for Csfr1b, are in direct contact with ovarian follicles, and thus are positioned to sense oocyte cues and activate ovarian macrophages. Analysis of recently published scRNA-seq data of human embryonic ovary (*37*) indicates the presence of *lhx9, csf1a*, and *irf8* expressing populations in the human early ovary (Fig. S4). To determine if Csf1a ligand was required for ovarian atresia and sex reversal, we generated *bmp15*;*csf1a DM*s. We found that removing Csf1a, the main ovarian-expressed ligand for Csf1rb, was sufficient to suppress ovarian failure and sex reversal of *bmp15* mutants (Fig. 4, A-F).

**Fig. 4.**
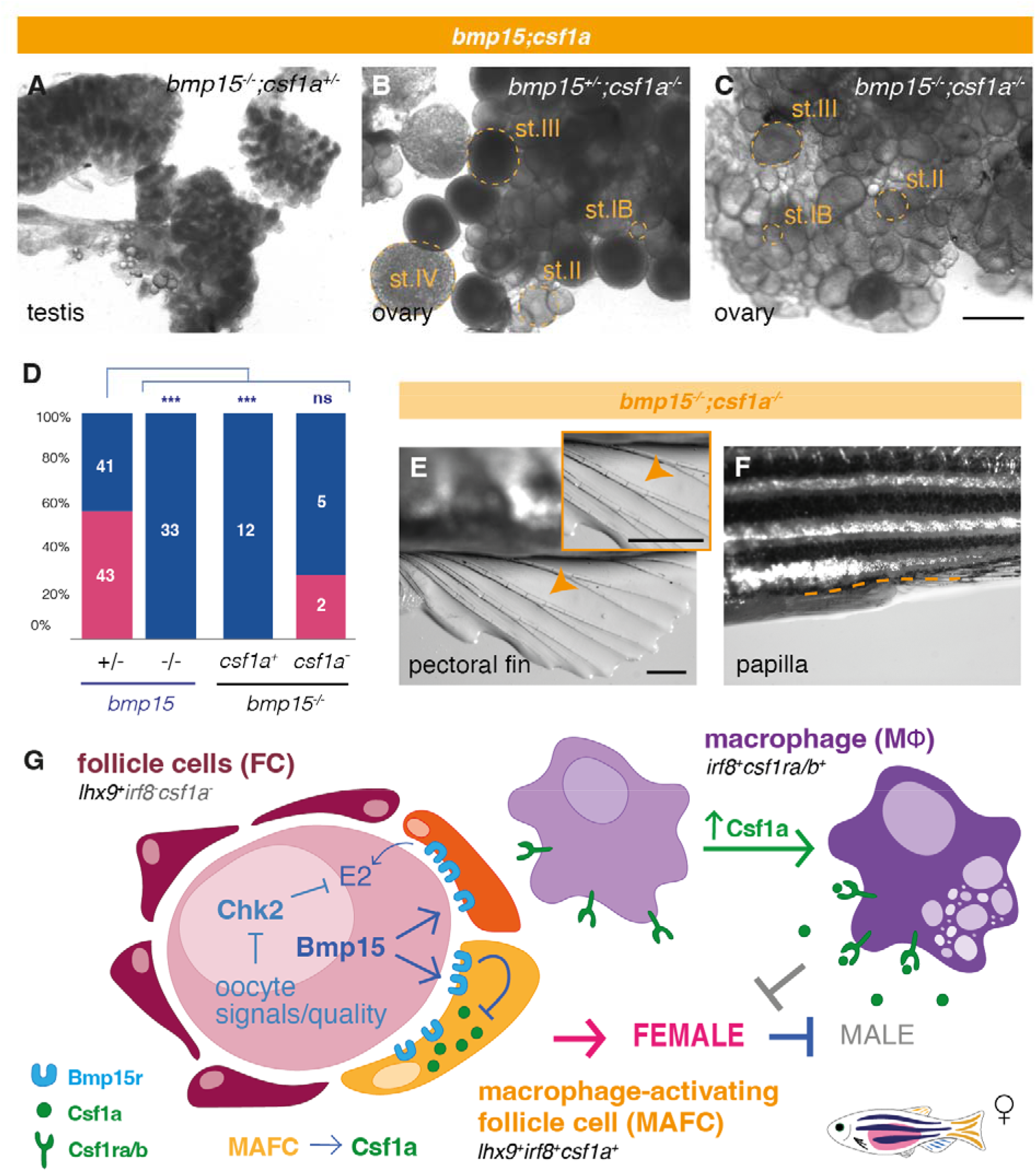
Csf1a activates macrophages and is required for ovarian failure and masculinization. **(A-C)** Live tissue images of adult *bmp15;csf1a* gonads of indicated genotypes. **(D)** Graph with sex ratios of indicated genotypes. Female (pink), male (blue). Numbers indicate individuals examined. Statistical analysis: Chi-square test with Bonferroni correction; p-Value comparisons to *bmp15*^+/-^, ***P ≤ 0.0001. **(E-F)** Images of *bmp15;csf1a* DM female secondary sex traits showing (E) absence of pectoral fin tubercles (magnified view in yellow box) and (F) female papilla. **(G)** Proposed model of germline-somatic-immune cell axis: signaling between oocyte derived Bmp15, follicle cells, and macrophages regulate ovarian follicle stability. Scale bars: (A-C) 500µm, (E, F) 1mm. St.I: stage I oocyte, St.II: stage II oocyte, St.III: stage III oocyte.

In zebrafish as in the mammalian ovary, BMP15 regulates both cell survival and cell fates (*3-5*). Therefore, we propose Bmp15 promotes follicle development and survival, and directly or indirectly (possibly via granulosa cells) silences MAFCs, which in response to failed follicle differentiation and oocyte loss release Csf1a ligand and signal to ovary macrophages to trigger sex reversal. Accordingly, MAFCs would act as specialized pre-follicle cells that express Csf1a, act as sentinels of oocyte quality, and activate macrophages to promote sex reversal when oocyte or granulosa signals decline (Fig. 4E).

Ovarian failure in humans has been associated with immunity-related disorders and genetic factors, like mutation of *bmp15*, but the specific immune cells and mechanisms driving premature follicle loss, infertility and masculinization were not known. This study shows that Bmp15 promotes follicle survival and cell fates required for follicle development, since *bmp15* mutant ovarian follicles preserved by loss of Chk2 could not progress developmentally. We discovered a novel population of ovarian prefollicle cells that express macrophage activating ligands. Further, we elucidate the molecular pathways and biological mechanisms underlying maintenance of ovarian follicles and identify definitive macrophages as key cell types activated by Csf1 ligand expressing sentinels that drive ovarian atresia and masculinization. Thus our work provides novel insights into the triggers of sex reversal in zebrafish and more broadly identifies potential cellular and molecular targets to prevent premature ovarian failure or ameliorate aspects of ovarian insufficiency/failure-related reproductive disorders.

## Acknowledgments

We thank members of the Marlow and Rangan labs for helpful discussions, The Center for Comparative Medicine staff at ISMMS for fish care, and the Microscopy CoRE at ISMMS.

## Author contributions

Conceptualization: PB, FLM

Methodology: PB, YL, BD, FLM

Investigation: PB, YL

Visualization: PB, YL, BD, FLM

Funding acquisition: FLM

Project administration: FLM

Supervision: FLM

Writing – original draft: PB, FLM

Writing – review & editing: PB, YL, BWD, FLM

## Data and materials availability

All data will be made freely available upon publication.

## Supplementary Materials

Materials and Methods

Figs. S1 to S4

Tables S1 to S2

References *(38-42)*

