## Supplemental materials for "Macrophage activation drives ovarian failure and masculinization"

### Macrophage activation elicits masculinization during ovarian failure.

**This PDF file includes:**

Materials and Methods
Figs. S1 to S4
Tables S1 to S2

### Materials and Methods

#### Animals

Mutant zebrafish lines used were: *bmp15*$^{uc31}$, *chk2*$^{sa20350}$, *csf1a*$^{re05}$, *csf1ra*$^{j1e4}$, *csf1rb*$^{re01}$, *irf8*$^{st96}$, *tp53*$^{zdf1}$, and Tg(*piwil1:egfp*)$^{uc02}$ *(26, 29, 32, 37-39)*. Experimental fish used for whole-mount immunohistochemistry and live tissue images were 3 months old or older (referred in figure legends as 'adults'). Fish used for scRNAseq libraries and single-molecule whole-mount RNA fluorescence *in situ* hybridization were 40 days old. Standard conditions were used for maintenance of all zebrafish. All protocols and procedures were performed following guidelines from the National Institutes of Health and approved by the Icahn School of Medicine at Mount Sinai Institutional (ISMMS) Animal Care and Use Committees (IACUC, 2017-0114).

#### Genotyping

Samples from fin clip, trunk or gonad were lysed in an alkaline lysis buffer (25 mM NaOH and 0.2 mM EDTA pH 12) to obtain genomic DNA (gDNA), heated at 95°C for 20 min, and cooled at 4°C before adding neutralization buffer (20 mM Tris-HCl and 0.1 mM EDTA pH 8.1) *(40)*. gDNA was PCR-amplified, followed when needed by restriction enzyme digestion and then resolved in a 3% 1:1 MetaPhor/agarose gel (primers and restriction enzymes used for each gene listed in Table S1).

#### Dissections

Fish were dissected after being anesthetized with a lethal dose of tricaine (MS-22) (400mg/l). All live tissue (without fixation) images of fish bodies were acquired with a Zeiss Stemi stereo microscope. Gonads were imaged using a Zeiss Zoom fluorescence stereo microscope.

Single-molecule whole-mount RNA *in situ* hybridization

Fish trunks were fixed overnight at 4°C in 4% paraformaldehyde and washed with PBS, dehydrated with MeOH, and then placed at −20°C overnight or until use. HCR RNA FISH probes were designed from Molecular Instruments (Table S2). Hybridization was performed following the manufacture's protocol (MI-Protocol-RNAFISH-Zebrafish) with the following changes: After rehydration in PBS with a series of MeOH>PBS 5 min washes, gonads were dissected and washed 4x 5 min in PBST (PBS + 1% Tween 20). Gonads were permeabilized with proteinase K at 50 µg/ml in PBST for 15 min. After the last manufacture's protocol step, samples were cleared in 1h-incubations of 30, 50, and 70% glycerol/PBS. Lastly, samples were mounted in ProLong Diamond Antifade Mountant with DAPI (Invitrogen) and imaged with a confocal microscope Leica SP8 STED (63X oil objective, 1024x1024 pixels format). All pictures were processed using ImageJ/FIJI.

Whole Mount Immunofluorescence

Tissues were fixed with 4% paraformaldehyde overnight at 4°C and washed the next day in PBS before dehydration with MeOH. Samples were stored at −20°C for at least one night or until use. Tissues were rehydrated with several washes of PBS before permeabilization with acetone for 10 minutes at −20°C and then incubated with 1) blocking buffer (5% normal goat serum/2% DMSO in 0.1% Tween/PBS) at room temperature for 1 hour or at 4°C overnight, 2) primary antibody overnight at 4°C, 3) washed in PBT, 4) incubated in secondary antibody at room temperature for 2 hours or overnight at 4°C 5) washed in PBT. To label germ cells, a Chicken anti-Vasa primary antibody *(41)* was used at a 1:3000 dilution (Table S2) followed by Alexa Fluor 488 secondary antibody (Molecular Probes) diluted 1:500. Whole mount tissues were mounted on slides using Vectashield with DAPI (Vector Laboratories) and imaged using a Zeiss Axio Observer inverted microscope equipped with Apotome.2 and a charged-coupled device (CCD) camera. All pictures were processed using ImageJ/FIJI and Adobe Illustrator.

Single-cell RNA sequencing (scRNAseq)

Single-cell RNA sequencing library expression data were from raw and processed data obtained from *(32)* for the zebrafish ovary, and from *(36)* for the fetal human ovary. UMAPS and graphs were generated using BBrowser3 software and online browsers embedded in the zebrafish (Single Cell Portal) and human (CellxCell) publications, following published analysis parameters.

**Fig. S1.**

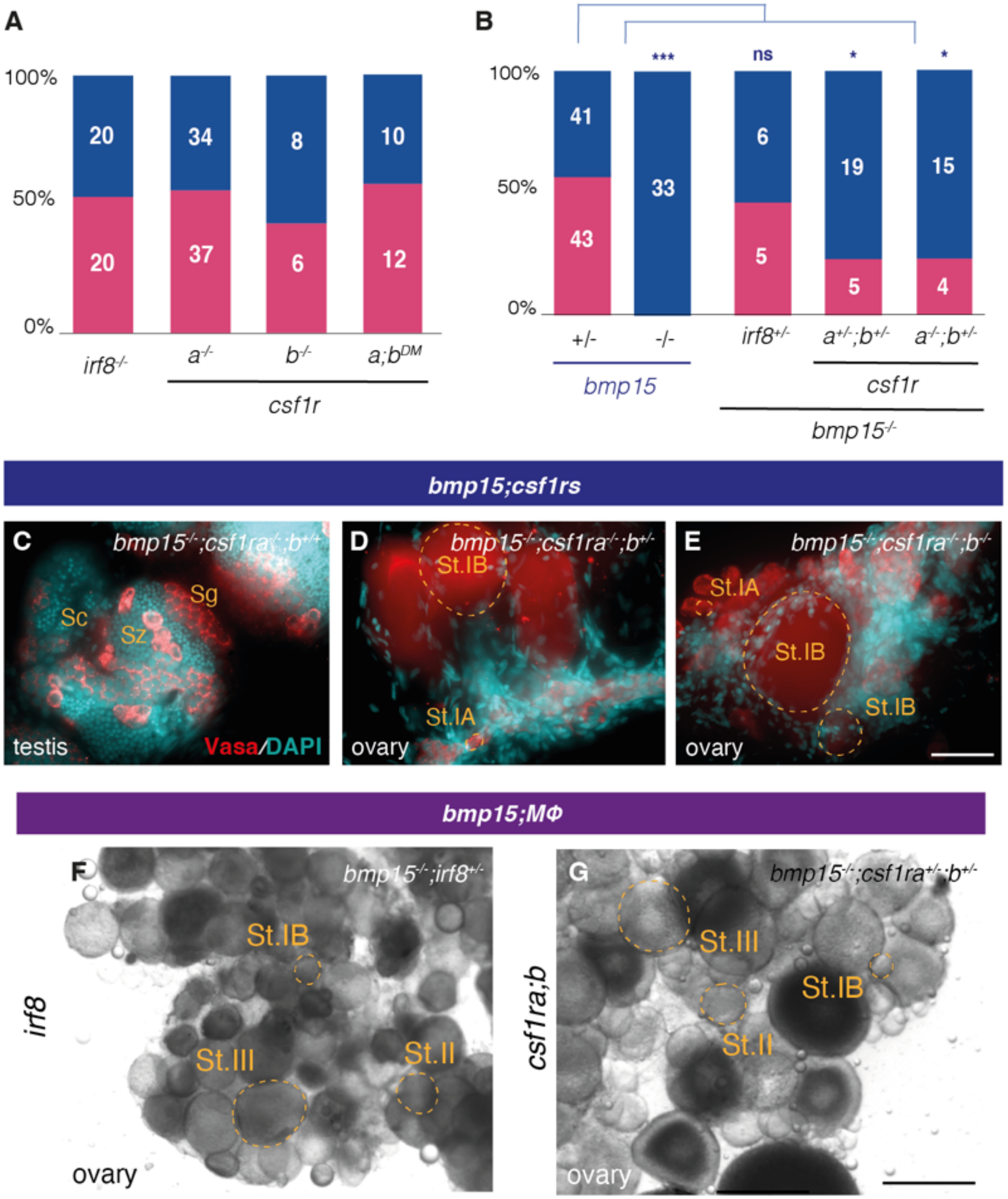


**Fig. S1. Absence of macrophages does not impar sex differentiation, but eliminating macrophages blocks ovarian failure and sex reversal of *bmp15* mutant females. (A-B)** Sex ratios graph of adult fish in the absence of (A) macrophages, or (B) macrophages and *bmp15*.

Female (pink); male (blue). Numbers indicate individuals examined. **(C-E)** Images of immunostained adult *bmp15* mutant gonad lacking (C) only primitive, (D) primitive and haploinsufficient for definitive, or (E) all macrophages. Vasa (red) labels germ cells. **(F-G)** Live tissue pictures of adult gonads of *bmp15* mutant fish heterozygous for (F) *irf8* or (G) *csf1rs*. Statistical analysis by Chi-square test with Bonferroni correction; p-Value comparisons are to $bmp15^{uc31/-}$, $^{*}P \leq 0.0125$, $^{***}P \leq 0.0001$. Scale bar: (C-E) 50µm, (F, G) 500µm. St.I: stage I oocyte, St.II: stage II oocyte, Sg: spermatogonia, Sc: spermatocyte, Sz: spermatozoa.

**Fig. S2.**

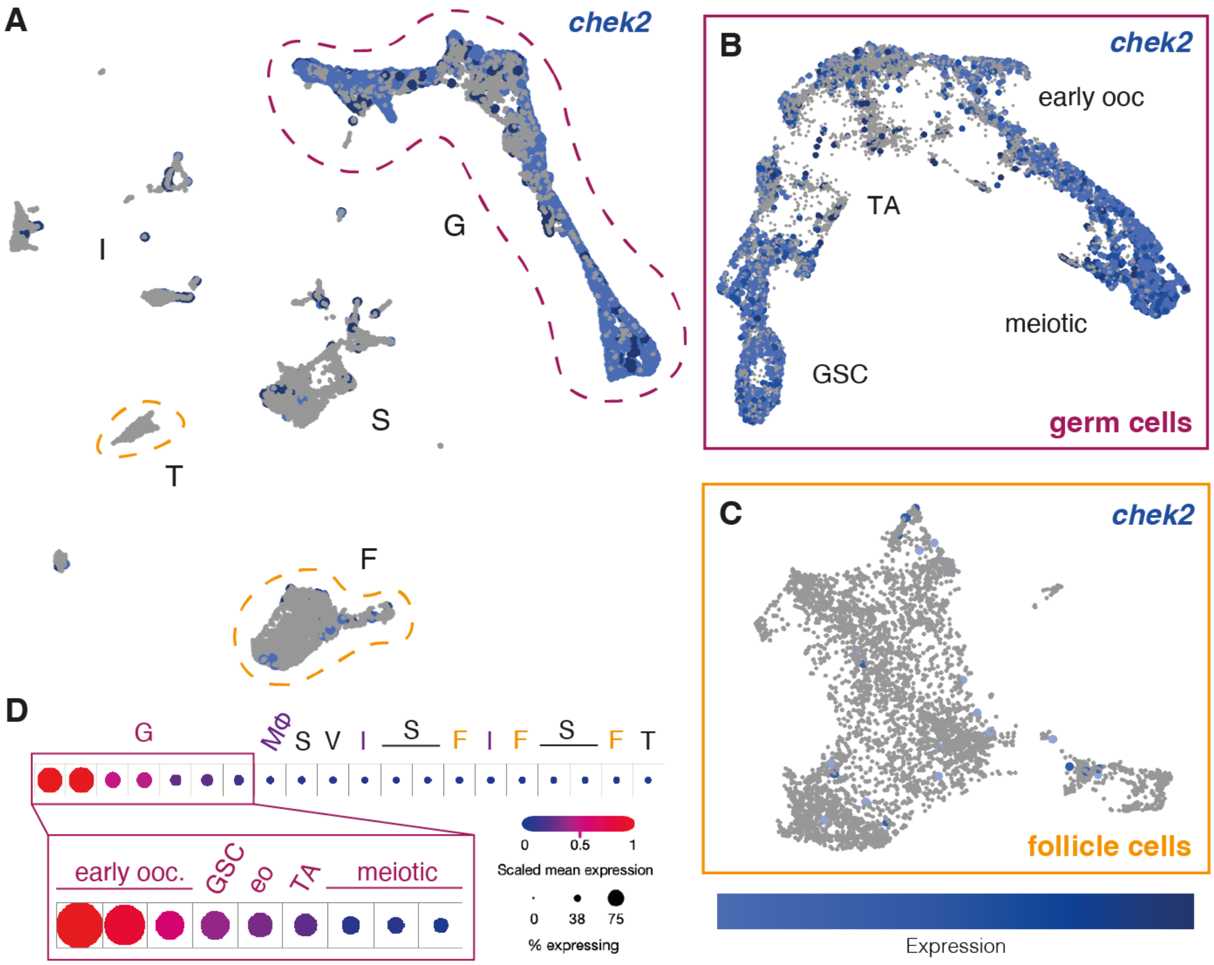


**Fig. S2. *chk2* is expressed in germline cells of the 40d ovary.** (**A-C**) UMAP plot of indicated gene expression in the 40d ovary in (A) all cells, (B) germ cells, and (C) follicle cells. (**D**) Analysis of *chek2* expression profile in specific clusters of cells within the ovary represented by Spearman's Rho correlation analysis. Expression in germ cells subcluster magnified in box. Early ooc: early oocyte, F: follicle, G: germline, GSC: germline stem cells, I: immune, Mϕ: macrophage, S: stroma, T: theca, TA: transit amplifying, V: vasculature.

**Fig. S3.**

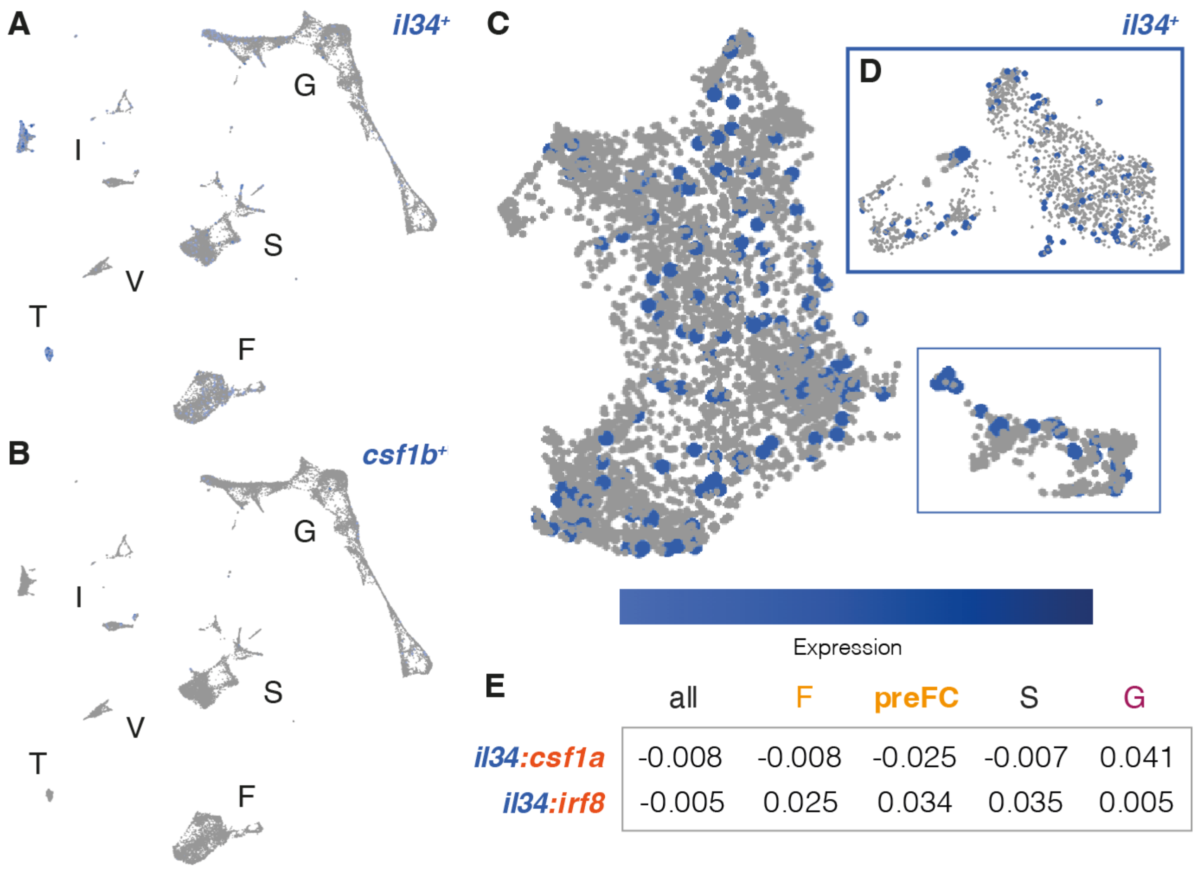


| | all | F | preFC | S | G |
|---|---|---|---|---|---|
| *il34:csf1a* | -0.008 | -0.008 | -0.025 | -0.007 | 0.041 |
| *il34:irf8* | -0.005 | 0.025 | 0.034 | 0.035 | 0.005 |

**Fig. S3. *csf1b* and *il34* Csf1R ligands are not expressed in pre-FCs. (A-D)** UMAP plot of indicated gene's expression in the 40dpf ovary in (A, B) all cells, (C) follicle cells, and (D) magnified subpopulation of pre-follicle cells. **(E)** Expression profiles of indicated genes in specific cluster of cells within the ovary represented by Spearman's Rho correlation analysis. F: follicle, G: germline, GSC: germ stem cells, I: immune, MΦ: macrophages, S: stroma, T: theca, V: vasculature.

Fig. S4.

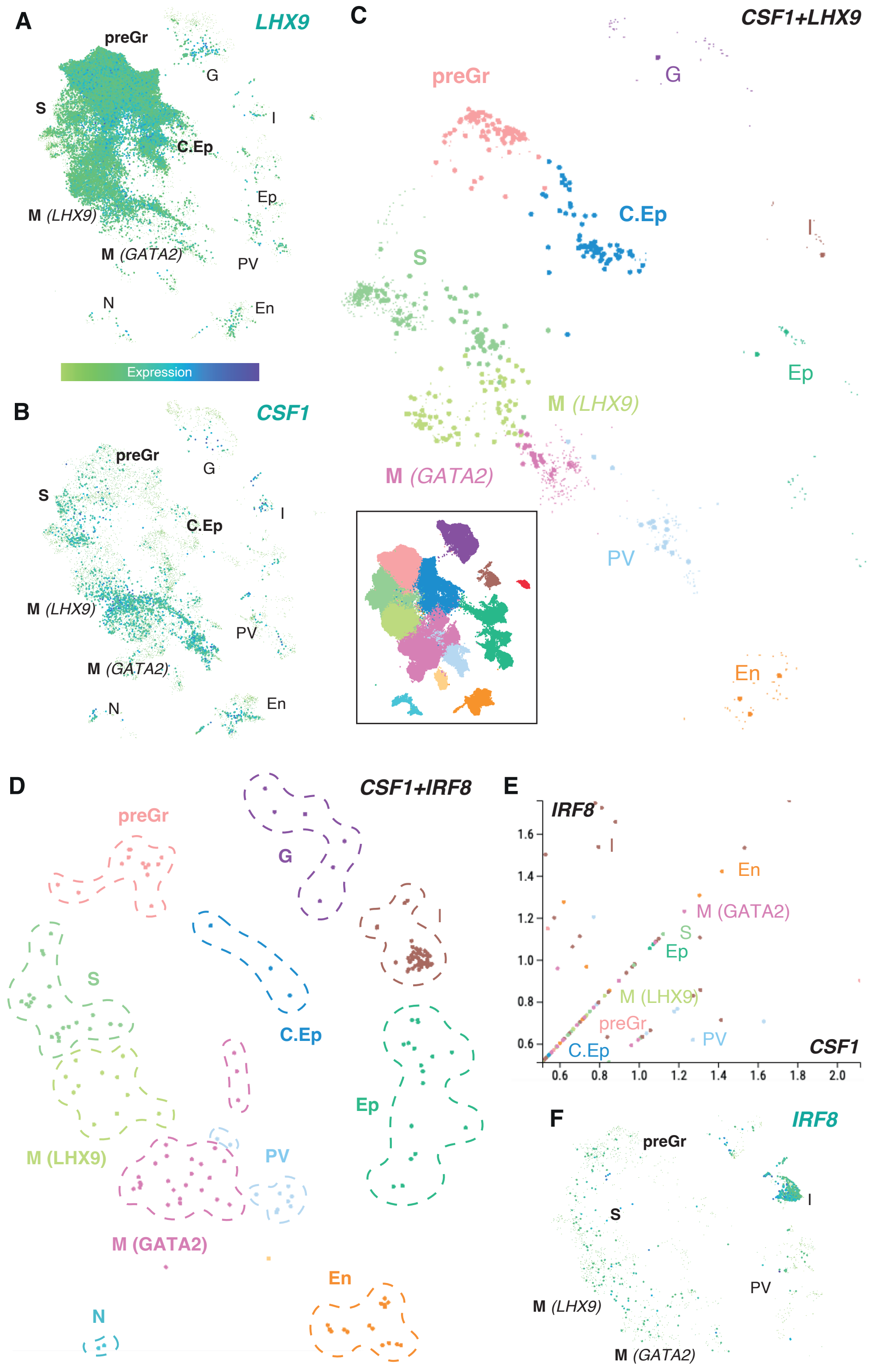


**Fig. S4. CSF1 source cells in human fetal ovary (A-D, F)** UMAP plots of indicated genes expression in the fetal human ovary showing (A, B, F) overall expression and (C, D) coexpression. UMAP clusters legend boxed in (C). (**E**) Scatterplot of cell cluster co-expressing indicated genes. C.Ep: coelomic epithelium, En: endothelial, Ep: epithelia, I: immune, M: mesenchymal, preGr, pregranulosa cells, PV: perivascular, S: supporting, N: neuro.

**Table S1.**

| Gene | Allele | F primer | R primer | RE (cuts) | line source |
|---|---|---|---|---|---|
| *bmp15* | ***uc31*** | AGCCTTTCAGGTGGCACTCG | CCACTGAAAAACACTTTCTCCC | - | Dranow et al., 2016 |
| *chek2* | ***sa20350*** | AGCCACACGAAATGCTGAG | CAGACTGAAGACTCCTACTACATTG | HpyCH4III (mut) | ZIRC (Derek Stemple) |
| *csf1a* | ***re05*** | GCCGGTTGAGCTTCTGAAAAT | GCATTTTGGTTAGGCTGCTG | - | Kuil et al., 2019 |
| *csf1ra* | ***j4e1*** | TCTGGGCAAAGAGGACAACATCACAC | CCACAGCTCTGCAAGGTTTG | SpeI (wt) | Parichy et al., 2000 |
| *csf1rb* | ***re01*** | GGACAGAGTTTTCGCTCCAG | ATTGGACTCCGCTCATGTTC | MspI (wt) | Oosterhof et al., 2018 |
| *irf8* | ***st96*** | ACATAAGGCGTAGAGATTGGACG | GAAACATAGTGCGGTCCTCATCC | AvaI (wt) | Shiau et al., 2015 |
| *tp53* | ***zdf1*** | ACATGAAATTGCCAGAGTATGTGTC | TCGGATAGCCTAGTGCGAGC | - | Berghmans et al., 2005 |

**Table S1. Experimental models and genotyping assays.**

**Table S2.**

| Name | [conc] | use | source (catalog #) |
|---|---|---|---|
| Chicken anti-zf Vasa | 1:3000 | IHC | Blokhina et al., 2019 |
| *csf1a* (amplifier B1) | 8nM | FISH | Molecular Instruments (custom) |
| *lhx9* (amplifier B3) | 8nM | FISH | Molecular Instruments (PRD333) |

**Table S2. Antibody and probes for fluorescent immunostainings and *in situ* hybridization.**
